# Differential effects of the second SARS-CoV-2 mRNA vaccine dose on T cell immunity in naïve and COVID-19 recovered individuals

**DOI:** 10.1101/2021.03.22.436441

**Authors:** Carmen Camara, Daniel Lozano-Ojalvo, Eduardo Lopez-Granados, Estela Paz-Artal, Marjorie Pion, Rafael Correa-Rocha, Alberto Ortiz, Marcos Lopez-Hoyos, Marta Erro Iribarren, Jose Portoles, Pilar Portoles, Mayte Perez-Olmeda, Jesus Oteo, Cecilia Berin, Ernesto Guccione, Antonio Bertoletti, Jordi Ochando

## Abstract

The rapid development and deployment of mRNA-based vaccines against the severe acute respiratory syndrome coronavirus-2 (SARS-CoV-2) led to the design of accelerated vaccination schedules that have been extremely effective in naïve individuals. While a two-dose immunization regimen with the BNT162b2 vaccine has been demonstrated to provide a 95% efficacy in naïve individuals, the effects of the second vaccine dose in individuals who have previously recovered from natural SARS-CoV-2 infection has been questioned. Here we characterized SARS-CoV-2 spike-specific humoral and cellular immunity in naïve and previously infected individuals during full BNT162b2 vaccination. Our results demonstrate that the second dose increases both the humoral and cellular immunity in naïve individuals. On the contrary, the second BNT162b2 vaccine dose results in a reduction of cellular immunity in COVID-19 recovered individuals, which suggests that a second dose, according to the current standard regimen of vaccination, may be not necessary in individuals previously infected with SARS-CoV-2.

## Manuscript

The BNT162b2 mRNA coronavirus disease 2019 (COVID-19) vaccine was authorized for emergency use by the Food and Drug Administration (FDA) and the European Medicines Agency (EMA) in December 2020 [1]. The vaccination strategy for the BNT162b2 vaccine involves an accelerated two-dose vaccination regimen administered 21 days apart, which has been demonstrated to induce a spike-specific humoral and cellular immunity associated with a 95% efficacy in naïve individuals [2]. However, in individuals with prior exposure to SARS-CoV-2, the utility of the second dose has been challenged. While robust spike-specific antibodies and T cells are induced by a single dose vaccination in SARS-CoV-2 seropositive individuals [3, 4], the second vaccination dose appears to exert a detrimental effect in the overall magnitude of the spike-specific humoral response in COVID-19 recovered individuals [5].

The effects of the second dose mRNA vaccine on spike-specific T cell expansion and contraction are however unknown in both naïve and in SARS-CoV-2 pre-exposed individuals. Understanding this gap of knowledge is critical since protection from disease severity and infection are likely to be dependent on the coordinated activation of both the humoral and cellular arms of adaptive immunity [6]. The complexity of monitoring virus-specific T cell magnitude and function has so far prevented the full characterization of the immune cellular response to the vaccine. To address this problem, we have shown that SARS-CoV-2-specific T cells (both CD4 helper and CD8 cytotoxic) can be reliably quantified in both SARS-CoV-2 infected individuals [7] and vaccine recipients [11] by a direct and quantitative ex vivo measurement of IFN-gamma secreted by whole blood samples stimulated with a pool of viral peptides.

Using this experimental approach, we studied the kinetics of T cell immunity at multiple time points (i.e. prior to, and 10 and 20 days after the first and second dose of vaccine) in 46 individuals with and without previous documented SARS-CoV-2 infection (23 naïve; mean age 39.9 years [range, 20 to 62], female 78%) and 23 PCR/antigen test-confirmed SARS-CoV-2 recovered (39% at 6-9 months after infection, 26% 3-6 months, 35% 1-3 months) (mean age 44.3 years [range, 20 to 76]; female 76%). Prior to vaccination, spike peptide pool-induced IFN-gamma production in whole blood of COVID-19 recovered individuals was higher than in naïve subjects (median pre-vaccination in naïve: 1.0 pg/ml [N=20] and in COVID-19 recovered: 46.9 pg/ml [N=21]) (**Fig.1A**). Using our modified protocol to evaluate IFN-gamma expression in peripheral blood mononuclear cells (PBMC) by flow cytometry [8], we demonstrated that SARS-CoV-2 spike and peptide pools stimulated the secretion of IFN-gamma in effector antigen-specific CD4^+^ effector T cells in COVID-19 recovered but not in naïve donors, confirming the presence of memory SARS-CoV-2 specific CD4^+^T cells in individuals with prior exposure to this virus (**Supp. Fig.1)** and that IFN-gamma released in whole blood reliably detects the Spike-specific T cell response.

**Fig. 1:**
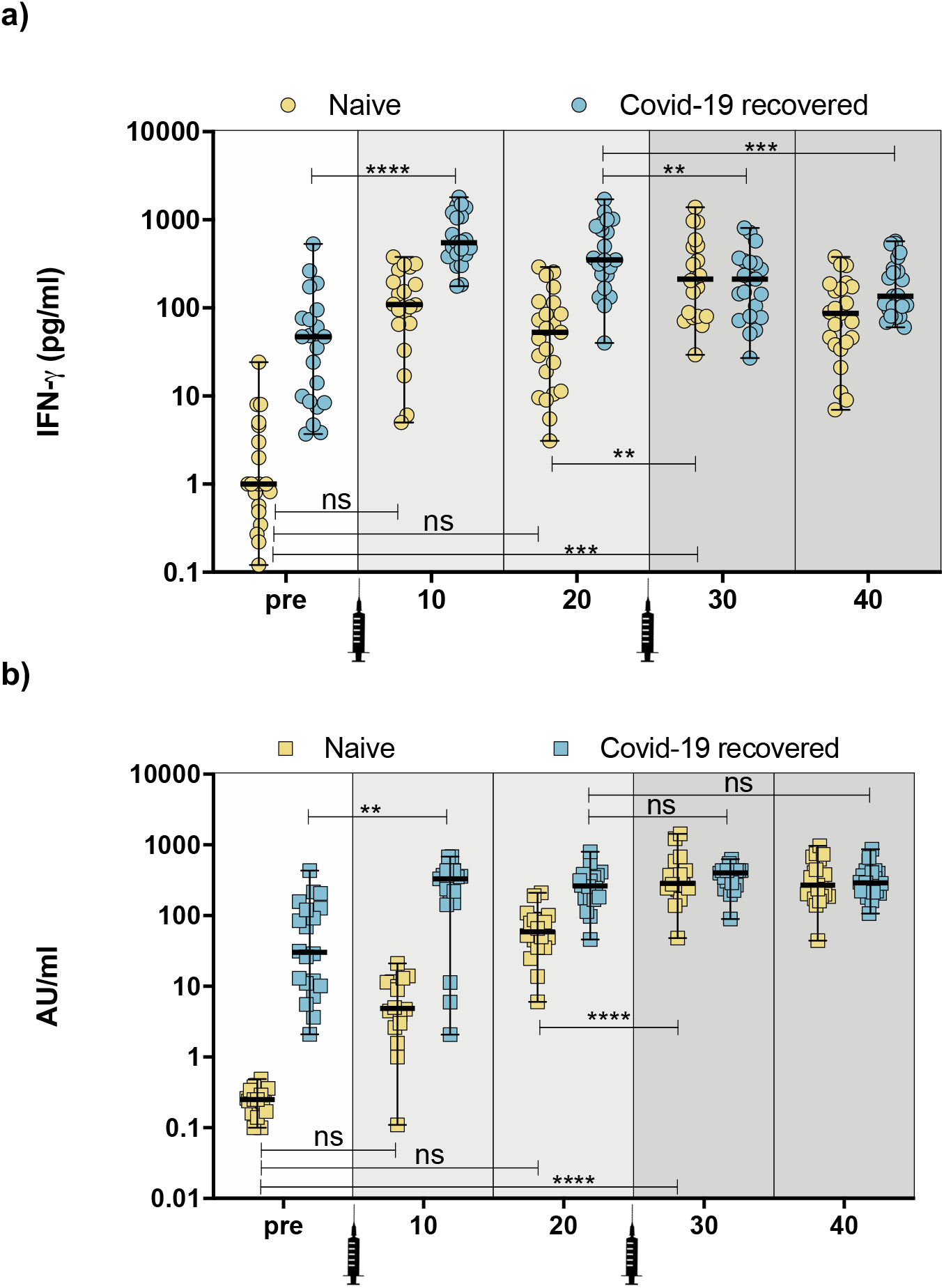
**a)** Quantification of IFN-gamma production in naïve and COVID-19 recovered individuals [pre-vaccination (pre), 10 and 20 days after the first vaccine dose (10 and 20, respectively), and 10 and 20 days after the second vaccine dose (30 and 40, respectively)] after overnight stimulation of whole blood with SARS-CoV-2 peptide pools. IFN-gamma concentration was determined using ELLA single plex cartridges. **b)** Kinetics of SARS-CoV-2 spike-specific IgG serum levels in naïve and COVID-19 recovered individuals measured by ACCESS SARS-CoV-2 CLIA. Statistical comparison between groups was performed using ANOVA test (** ≤ 0.003; *** ≤ 0.0003; *** ≤ 0.0001).

Evaluation of the spike-specific T cell response 10 days after the first dose indicates that COVID-19 recovered individuals mount a stronger IFN-gamma response in comparison with naïve subjects (median day 10 post-first vaccine dose in naïve: 110.4 pg/ml [N=20] and in COVID-19 recovered: 520 pg/ml [N=21]) (**Fig.1A**). Interestingly, while COVID-19 recovered individuals maintain their T cell immunity on day 20 post-first vaccine dose, IFN-gamma response in naïve individuals rapidly decreases (median day 20 post-first vaccine dose in naïve: 31.3 pg/ml [N=23] and in COVID-19 recovered: 278.0 pg/ml [N=21]). These results indicate that individuals with pre-existing immunity exert a more potent and sustained T cell response to SARS-CoV-2 spike after the first dose of the vaccine, consistent with recent investigations [12].

We next studied the effects of the second dose of the vaccine. Sampling on day 10 after the second dose confirmed the beneficial effects of the recall vaccine in naïve individuals who increase their IFN-gamma to significant levels (median day 10 post-second vaccine dose in naïve: 162.0 pg/ml [N=20] vs median pre-vaccination in naïve: 1.0 pg/ml [N=20]; p=0.0003). Interestingly, we observed a similar trend when measuring SARS-CoV-2 spike-specific IgG levels, which indicates that naïve individuals achieve significant concentrations of IgG antibodies after the second vaccine dose (**Fig.1B**), suggesting that protection is achieved following the standard two-dose regimen for COVID-19 vaccination in naïve individuals.

Unexpectedly, COVID-19 recovered individuals significantly decreased their IFN-gamma production on day 10 after the second vaccine dose (median day 10 post-second vaccine dose in COVID-19 recovered: 212.0 pg/ml [N=21] vs median pre-vaccination in COVID-19 recovered: 46.9 pg/ml [N=21]; p=0.0032) (**Fig.1A**). These findings indicate that, while naïve subjects significantly increase their immunity against SARS-CoV-2 spike protein after the second dose of the vaccine, COVID-19 recovered individuals do not seem to benefit from the standard regimen for COVID-19 vaccination. We observed no differences in IFN-gamma secretion between naïve and COVID-19 recovered individuals on day 20 after the second dose (median day 20 post-second vaccine dose naïve: 87.5 pg/ml [N=23] and COVID-19 recovered: 136.0 pg/ml [N=23]).

Our analysis of humoral and cellular spike-specific immunity in recipients of the BNT162b2 mRNA COVID-19 vaccine indicates that a single dose of the vaccine elicits the generation of specific antibodies and a T cell immune response in naïve and COVID-19 recovered individuals, consistent with recently published data [4] and [12]. In naïve subjects, the T cell response decays rapidly after the first vaccine dose, but is boosted in conjunction with spike-specific IgG levels after the second dose. On the contrary, in individuals with a pre-existing immunity against SARS-CoV-2, the second vaccine dose not only fail to boost humoral immunity (**Fig.1B**) but determines a contraction of the spike-specific T cell response. Thus, these data argue in favor of meeting the vaccination scheme determined in clinical trials with prompt administration of the second dose in individuals without previous SARS-CoV-2 exposure [9]. On the other hand, individuals with pre-existing immunity against SARS-CoV-2 should be spared the second dose of the vaccine, at least temporarily, to prevent a possible contraction of their spike-specific memory T cell immunity. Our study has clear limitations.

Due to its observational nature, the mechanisms of the contraction of the spike-induced production of IFN-gamma in COVID-19 recovered subjects observed after the second vaccination dose were not investigated. We can only hypothesize that the effector memory CD4^+^ T cells expanded by first vaccine dose in COVID-19 recovered individuals may be prone to activation-induced cell death (AICD) after the second vaccination dose. We cannot however exclude that the second dose of the vaccine may only functionally exhaust the spike-specific T cells without really reducing the quantity of the long-term pool of effector memory T cells. Therefore, more detailed analysis of the phenotype of the spike-specific T cells induced by COVID-19 vaccines both in naïve and recovered individuals are needed to answer these questions.

At present, there is a shortage of doses to vaccinate a large proportion of the population and public health authorities are designing priority vaccination strategies according to age, comorbidities and risk assessment criteria. We conclude that using *ex vivo* SARS-CoV2 specific cellular immunity tests may allow health authorities to screen individuals for pre-existing immunity against SARS-CoV-2 and update their priority strategy for ongoing vaccination campaigns against COVID-19 to provide vaccination certificates to those individuals with pre-existing immunity after the first vaccine dose.

## Supporting Figure 1

**Supp. Fig.1:**
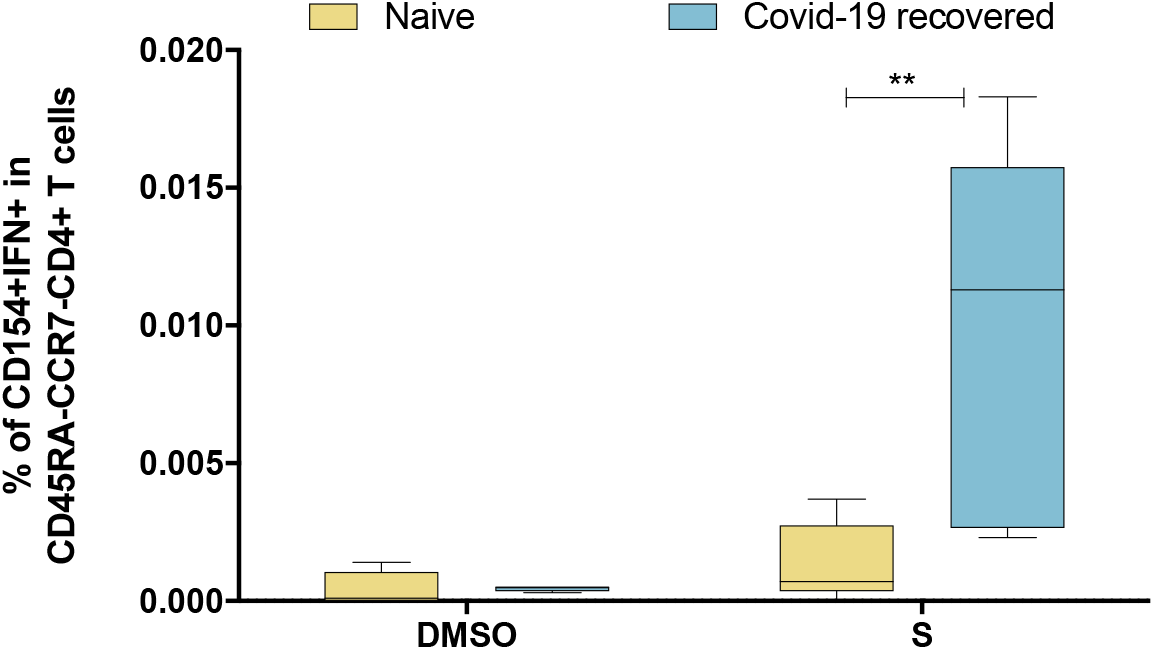
Flow cytometry determination of IFN-gamma-producing antigen-specific (CD154^+^) memory (CD45RA^−^CCR7^−^) CD4^+^ T cells in peripheral blood mononuclear cells obtained from naïve and COVID-19 recovered individuals (6 months after PCR confirmation) unstimulated (DMSO) or stimulated with SARS-CoV-2 spike (S) peptide pools. Statistical comparison between groups was performed using t-test (** ≤ 0.003).

## Materials and Methods

### SARS-CoV-2 Peptide pools

SARS-CoV-2 peptide pools of 15-mers (55 peptides) covering 40.5% of the Spike protein and contain most of the SARS-CoV2 spike epitope published to date [10] were used as reported in [7]

### Whole blood culture with SARS-CoV-2 peptide pools

320 μl of whole blood drawn on the same day were mixed with 80 μl RPMI and stimulated with pools of SARS-CoV-2 peptides (S; 2 μg/ml) or a DMSO control. After 15 hours of culture, the supernatant (plasma) was collected and stored at −80°C until quantification of cytokines.

### Cytokine quantification and analysis

Cytokine concentrations in the plasma were quantified using Ella with microfluidic multiplex cartridges measuring IFN-γ following the manufacturer’s instructions (ProteinSimple, San Jose, California). The level of cytokines present in the plasma of DMSO controls was subtracted from the corresponding peptide pool stimulated samples.

### Spike-specific IgG quantification

The ACCESS SARS-CoV-2 CLIA (Beckman Coulter Inc., California, USA) was used for semiquantitative detection of IgG directed against S protein RBD using serum obtained from venipuncture blood. Samples were tested on a UniCel Dxl 800 high-performance analyser.

### Flow Cytometry

Subjects were recruited from the Mount Sinai School of Medicine under an institutional review board–approved protocol and informed consent obtained from all subjects. Peripheral blood mononuclear cells (PBMCs) were isolated by means of density centrifugation with Ficoll-Paque Plus (GE Healthcare, Pittsburgh, PA) and cultured in AIM-V medium (Gibco, Grand Island, NY) with 2.5% AB human serum (Gemini Bio-Products, Inc., West Sacramento, CA). PBMCs were unstimulated (DMSO) or stimulated with SARS-CoV-2 spike (S) peptide pools (5 mg/mL) in presence of GolgiPlug (BD Biosciences, San Jose, CA) for 6 hours. Harvested PBMCs were stained for viability (Live/Dead Fixable, Invitrogen, Carlsbad, CA), washed and stained for surface markers, fixed in 4% paraformaldehyde (Electron Microscopy Services, Hatfield, PA), and treated with permeabilization buffer (eBioscience, San Diego, CA) before staining with labeled antibodies to detect intracellular CD154 and cytokines. Stained cells were analyzed subsequently on a CytekTM Aurora device (Cytek Biosciences, Fremont, CA).

### Statistical analysis

For IgG and IGN-gamma determination, statistical comparison between groups was performed using ANOVA test (** ≤ 0.003; *** ≤ 0.0003; *** ≤ 0.0001). For flow cytometry studies, statistical comparison between groups was performed using t-test (** ≤ 0.003).

## Author contributions

Overall design of the project: EG, AB and JO. Acquisition of experimental data: All co-authors. Generation of reagents and scientific inputs: All co-authors. JO, EG and AB wrote the manuscript with input from all co-authors.

## Acknowledgments

We acknowledge the technical contribution of Arroyo Sánchez, Daniel; Baranda, Jana; Baztan-Morales, Sara; Bodega-Mayor, Irene; Castillo de la Osa, María; Cervera, Isabel; Comins-Boo, Alejandra; del Álamo Mayo, Carmen; Gil-Manso, Sergio; Gonzalez Beatriz; Gonzalez-Perez, Maria; Hatem, Sandra; Irure-Ventura, Juan; Miguens, Iria; Montes-Casado, Maria; Muñoz Martinez, Sara; Ojeda, Gloria; Pereira, Monica; Rodrigues-Guerreiro, Catarina; Rodriguez-Garcia, Mercedes; Rojo-Portolés, Maria Pilar; San Segundo, David; Sanchez-Tarjuelo, Rodrigo and Schwarz, Megan. We would also like to acknowledge Beckman Coulter for donating the equipment required for the determination of spike-specific IgG antibodies. This work was supported by ISMMS seed fund to EG and JO; Instituto de Salud Carlos III, COV20-00668 to RCR; Instituto de Salud Carlos III, Spanish Ministry of Science and Innovation (COVID-19 Research Call COV20/00181) co-financed by European Development Regional Fund “A way to achieve Europe” to EP; Instituto de Salud Carlos III, Spain (COV20/00170); Government of Cantabria, Spain (2020UIC22-PUB-0019) to MLH; Instituto de Salud Carlos III (PI16CIII/00012) to PP; Fondo Social Europeo e Iniciativa de Empleo Juvenil YEI (Grant PEJ2018-004557-A) to MPE; REDInREN 016/009/009 ISCIII; This project has received funding from the European Union’s Horizon 2020 research and innovation programme VACCELERATE under grant agreement No [101037867] to JO.

## Ethics Statement

The study protocols for the collection of clinical specimens from individuals with and without SARS-CoV-2 infection were reviewed and approved by Hospital La Paz, Hospital 12 de Octubre, Hospital Gregorio Marañón, IIS-Fundación Jimenez Díaz, Hospital Universitario Marqués de Valdecilla-IDIVAL and Hospital Puerta de Hierro Clinical Research Ethics Committee (CEIm), and Mount Sinai Hospital Institutional Review Board (IRB).

## References

1. Krammer, F., SARS-CoV-2 vaccines in development. Nature, 2020. 586(7830): p. 516–527.

2. Polack, F.P., et al., Safety and Efficacy of the BNT162b2 mRNA Covid-19 Vaccine. N Engl J Med, 2020. 383(27): p. 2603–2615.

3. Krammer, F., et al., Antibody Responses in Seropositive Persons after a Single Dose of SARS-CoV-2 mRNA Vaccine. N Engl J Med, 2021.

4. Prendecki, M., et al., Effect of previous SARS-CoV-2 infection on humoral and T-cell responses to single-dose BNT162b2 vaccine. Lancet, 2021.

5. Samanovic, M.I., et al., Poor antigen-specific responses to the second BNT162b2 mRNA vaccine dose in SARS-CoV-2-experienced individuals. medRxiv, 2021.

6. McMahan, K., et al., Correlates of protection against SARS-CoV-2 in rhesus macaques. Nature, 2021. 590(7847): p. 630–634.

7. Le Bert, N., et al., Highly functional virus-specific cellular immune response in asymptomatic SARS-CoV-2 infection. J Exp Med, 2021. 218(5).

8. Tan, A.T., et al., Early induction of functional SARS-CoV-2-specific T cells associates with rapid viral clearance and mild disease in COVID-19 patients. Cell Rep, 2021. 34(6): p. 108728.

9. Kadire, S.R., R.M. Wachter, and N. Lurie, Delayed Second Dose versus Standard Regimen for Covid-19 Vaccination. N Engl J Med, 2021. 384(9): p. e28.

10. Grifoni, A., et al., A Sequence Homology and Bioinformatic Approach Can Predict Candidate Targets for Immune Responses to SARS-CoV-2. Cell Host Microbe, 2020. 27(4): p. 671–680 e2.

11. Kalimudin, S., et al., Early T Cell and Binding Antibody Responses are Associated with COVID-19 RNA Vaccine Efficacy Onset. http://dx.doi.org/10.2139/ssrn.3796533.

12. Tauzin, A., et al., A single BNT162b2 mRNA dose elicits antibodies with Fc-mediated effector functions and boost pre-existing humoral and T cell responses. https://doi.org/10.1101/2021.03.18.435972

